# Genomic epidemiology and global diversity of the emerging bacterial pathogen *Elizabethkingia anophelis*

**DOI:** 10.1101/044792

**Authors:** Sebastien Breurec, Alexis Criscuolo, Laure Diancourt, Olaya Rendueles, Mathias Vandenbogaert, Virginie Passet, Valérie Caro, Eduardo P. C. Rocha, Marie Touchon, Sylvain Brisse

**Affiliations:** Unité Environnement et Santé, Institut Pasteur de Guadeloupe, Pointe-à-Pitre, France, Faculté de Médecine Hyacinthe Bastaraud, Université des Antilles, Pointe-à-Pitre, France, Laboratoire de Microbiologie clinique et environnementale, Centre Hospitalier Universitaire de Pointe-à-Pitre/les Abymes, Pointe-à-Pitre, France; Laboratoire de Microbiologie Clinique, Institut Pasteur de Bangui, Bangui, Central African Republic; Institut Pasteur, Hub Bioinformatique et Biostatistique, C3BI, USR 3756 IP CNRS, Paris, France; Unité Environnement et Risques Infectieux, Institut Pasteur, Paris, France; Microbial Evolutionary Genomics, Institut Pasteur, Paris, France; UMR 3525, CNRS, Paris, France

## Abstract

*Elizabethkingia anophelis* is an emerging pathogen. Genomic analysis of strains from clinical, environmental or mosquito sources is needed to understand the epidemiological emergence of *E. anophelis* and to uncover genetic elements implicated in antimicrobial resistance, pathogenesis, or niche adaptation. Here, the genomic sequences of two nosocomial isolates that caused neonatal meningitis in Bangui, Central African Republic, were determined and compared with *Elizabethkingia* isolates from other world regions and sources. Average nucleotide identity firmly confirmed that *E. anophelis, E. meningoseptica* and *E. miricola* represent distinct genomic species and led to re-identification of several strains. Phylogenetic analysis of *E. anophelis* strains revealed several sublineages and demonstrated a single evolutionary origin of African clinical isolates, which carry unique antimicrobial resistance genes acquired by horizontal transfer. The *Elizabethkingia* genus and the species *E. anophelis* had pan-genomes comprising respectively 7,801 and 6,880 gene families, underlining their genomic heterogeneity. African isolates were capsulated and carried a distinctive capsular polysaccharide synthesis cluster. A core-genome multilocus sequence typing scheme applicable to all *Elizabethkingia* isolates was developed, made publicly available (http://bigsdb.web.pasteur.fr/elizabethkingia), and shown to provide useful insights into *E. anophelis* epidemiology. Furthermore, a clustered regularly interspaced short palindromic repeats (CRISPR) locus was uncovered in *E. meningoseptica, E. miricola* and in a few *E. anophelis* strains. CRISPR spacer variation was observed between the African isolates, illustrating the value of CRISPR for strain subtyping. This work demonstrates the dynamic evolution of *E. anophelis* genomes and provides innovative tools for *Elizabethkingia* identification, population biology and epidemiology.

**IMPORTANCE:** *Elizabethkingia anophelis* is a recently recognized bacterial species involved in human infections and outbreaks in distinct world regions. Using whole-genome sequencing, we showed that the species comprises several sublineages, which differ markedly in their genomic features associated with antibiotic resistance and host-pathogen interactions. Further, we have devised high-resolution strain subtyping strategies and provide an open genomic sequence analysis tool, facilitating the investigation of outbreaks and tracking of strains across time and space. We illustrate the power of these tools by showing that two African healthcare-associated meningitis cases observed 5 years apart were caused by the same strain, providing evidence that *E. anophelis* can persist in the hospital environment.

## INTRODUCTION

*Elizabethkingia* is a genus of aerobic, non-motile, Gram-negative rods belonging to family *Flavobacteriaceae* in the phylum *Bacteroidetes. Elizabethkingia* strains are considered ubiquitous in nature, and the genus currently comprises three species. *E. meningoseptica* is a common cause of neonatal meningitis, especially in premature infants during the first weeks of life, and also causes a wide range of human infections as an opportunistic pathogen (1-3). *E. miricola* is rarely associated with sepsis (4). *E. anophelis* was recently described from the midgut of *Anophelis gambiae* mosquitoes (5). Shortly after, some of us described the two first human cases of neonatal meningitis, caused by *E. anophelis* in Bangui, Central African Republic (6). A nosocomial origin was suspected for these strains given that patients had a history of intubation and mechanical ventilation at birth. The two meningitis cases occurred in the same hospital five years apart and an epidemiological link between them was not established. *E. anophelis* was also implicated in an outbreak in an intensive care unit in Singapore (7) and in a transmission from mother to infant in Hong Kong (8), resulting in late-onset meningitis in the baby. At the time of writing, an unprecedented large outbreak of E. anophelis from a yet unknown source is ongoing in Wisconsin, USA. *E. anophelis* is therefore considered as an emerging opportunistic pathogen. In all previously reported clinical cases (6-8), *E. anophelis* isolates were resistant to most antibiotics recommended for empirical therapy, including third-generation cephalosporins, aminoglycosides and carbapenems (9), probably accounting for the poor infection outcomes. Consistently, most *E. anophelis* genomes harbor multiple antimicrobial resistance genes, including the two metallo-betalactamase genes *bla*_GOB_ and *bla*_blaB_, associated with resistance to carbapenems (10). Currently, the clinical significance and biological distribution of *E. anophelis* is restricted, as this novel species is not well distinguished from other *Elizabethkingia* species in routine microbiological identification procedures, and as reference identification of *E. anophelis* relies mostly on the low-resolution 16S rRNA gene.

Genomic studies of *E. anophelis* strains from clinical, environmental and mosquito origins have revealed interesting features that may contribute to the adaptation of this bacterium to mosquito and that may be implicated in its pathogenesis (8, 11, 12). Genomic sequencing has also been used to derive epidemiological insights during hospital outbreaks (8, 12). However, knowledge on the genomic diversity and population structure of *E. anophelis* is still scant, and there is currently no standard way to compare isolates at local, regional or global scales, which limits our understanding of the epidemiology and global emergence of *E. anophelis*.

The aims of this study were to determine the phylogenetic origin and genomic features of the two African neonatal meningitis *E. anophelis* isolates; to compare them with *Elizabethkingia* isolates from other world regions and sources; to take advantage of the high resolution of whole genome sequencing to distinguish *Elizabethkingia* species unambiguously, and to develop a publicly-accessible genotyping system enabling a common language on strain types in order to ease future collaborations on biodiversity studies and epidemiological investigations of *Elizabethkingia* clinical and environmental isolates.

## RESULTS AND DISCUSSION

### Genome sequences of African isolates E27017 and E18064

The two African isolates were sequenced using paired-end Illumina technology. The genome sequence of isolate E18064 was assembled into 213 contigs of total length 4,080,763 bp. It is 35.7% G+C rich and contains 3,648 CDSs. The E27107 genome sequence is 4,059,474 bp long, 35.5% G+C rich and contains 3,674 CDSs distributed over 89 contigs. The genome characteristics of the two African isolates thus appeared similar to those of previously sequenced *Elizabethkingia* genomes (Table S1).

### Genome-based identification of members of the *Elizabethkingia* genus

To determine with confidence the species-level identification of the two African and the 18 other *Elizabethkingia* isolates, we calculated average nucleotide identity (ANI) based on their genomic sequences (Table S2). The distribution of ANI values among the 20 genomes classified them into three main groups (Fig. 1a) and showed a clear separation of *E. anophelis* strains from *E. meningoseptica* and *E. miricola*. These results fully demonstrate that these three taxa represent distinct genomic groups. The two African isolates clearly belong to *E. anophelis* based on their high ANI with R26^T^, the taxonomic type strain of this species, fully confirming their initial identification (6). Even though they were initially identified as *E. meningoseptica*, the three strains 502, Endophthalmitis and B2D also appeared closely related to *E. anophelis* isolates based on ANI values. Therefore, these three strains should be re-assigned to *E. anophelis*. Based on its ANI value with *E. miricola* reference strain ATCC 33958, the unclassified *Elizabethkingia* sp. strain BM10 isolated from a termite, clearly belongs to *E. miricola*. Urea degradation is a differential phenotypic characteristic of *E. miricola*, and we confirmed the presence in both *E. miricola* genomes of the urease cluster, whereas it was absent from all other genomes. The ANI values among *E. anophelis* genomes ranged from 96.97% to 100% (Fig. 1b), thus clearly higher than the classical 95% threshold used for species delineation. This result confirms that *E. anophelis* isolates belong to a single genomic species. Of note, the ANI value estimated between *E. anophelis* and *E. meningoseptica* was only 79.7%, showing that these species are separated by an evolutionary distance similar to the one observed between *Escherichia coli* and *Salmonella* spp. (80.4%), two model species with very distinct ecological lifestyles and virulence features.

**FIG 1.**
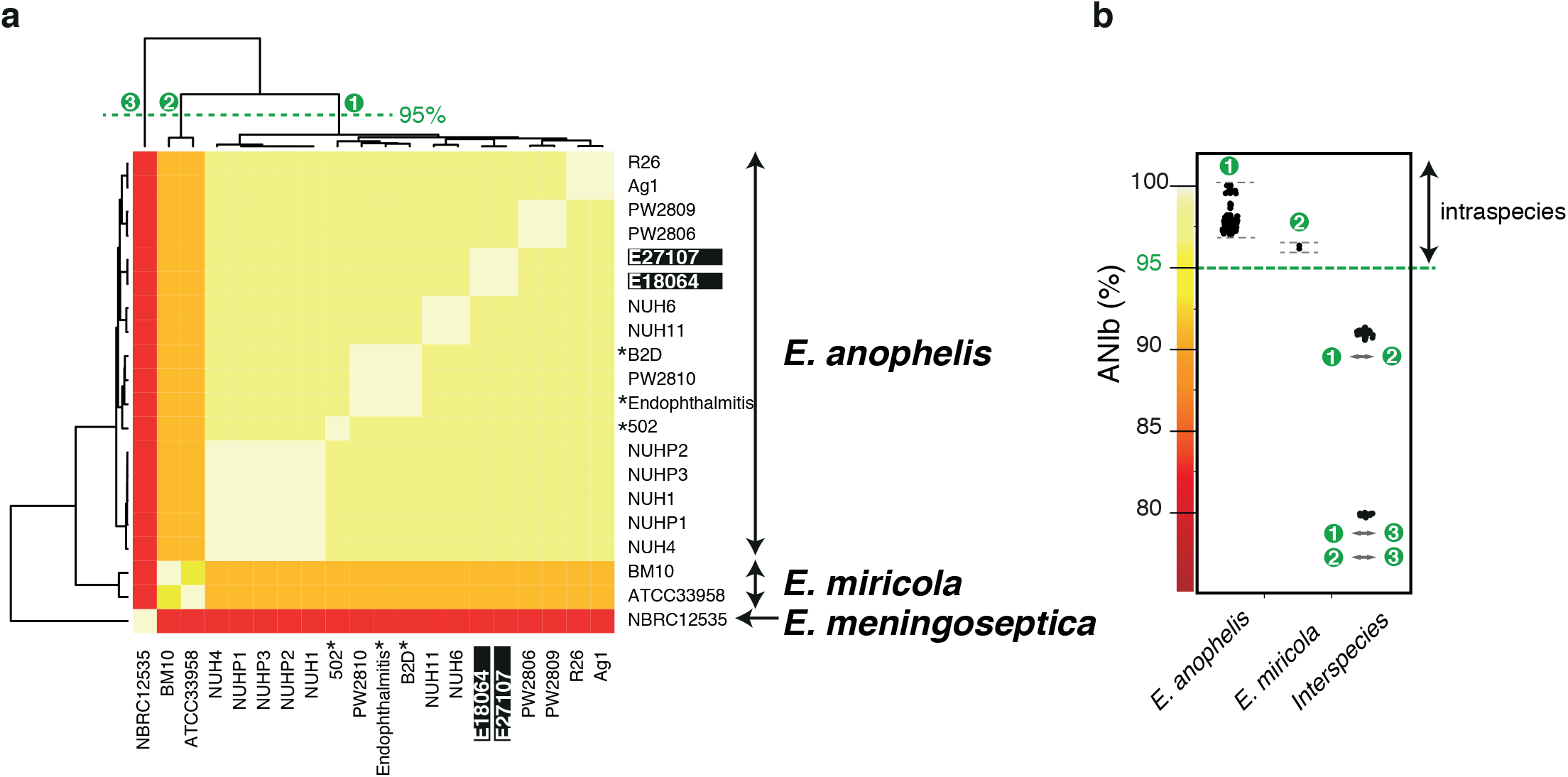
Heat-map of the ANIb for each pairwise comparison (a) and intra and inter-species ANIb variability (b). Green dashed line on the top dendogram marks groups of taxa that are from the same species or have more than 95% ANIb values and therefore might be regarded as part of the same species. Black asterisks indicate strains previously described as *E. meningoseptica*. Black rectangles indicate the two hospital-acquired isolates from Central African Republic sequenced in this study.

The two African *E. anophelis* isolates were initially identified based on 16S rRNA sequencing (6). Phylogenetic analysis of the 16S rRNA sequences (Fig. S1a) was consistent with the clustering into three species. However, four *E. anophelis* isolates (502, B2D, Endophthalmitis and PW2810) were linked to the other *E. anophelis* strains in a clade with an atypically long branch. Inspection of the alignment revealed that this long branch was caused by 10 nucleotide differences clustered within a single region of 35 nucleotides (Fig. S1b). These results indicate that careful interpretation may be required when classifying *Elizabethkingia* isolates based on the divergence values estimated from the classical 16S rRNA marker.

### Gene repertoire of *Elizabethkingia*

To compare the gene repertoire of the two African isolates with other *Elizabethkingia* isolates, we first quantified the diversity of *Elizabethkingia* genes by computing the set of ubiquitous genes (core genome) and the set of different homologous genes families (pangenome) among the 20 genomes. At the level of the genus *Elizabethkingia*, the core genome contained 2,221 orthologous protein families, corresponding to 65% of the size of the smallest proteome observed for *E. meningoseptica* NBRC 12535. The core-genome of the species *E. anophelis* had 2,512 orthologous protein families, corresponding to 72% of the size of the smallest proteome (strain B2D). Gene rarefaction analyses showed that the *E. anophelis* core genome varies little with the addition of the last genomes (Fig. 2a), suggesting that this core genome estimate is robust. Both the pan-genomes of the *Elizabethkingia* genus and of *E. anophelis* were large with, respectively, 7,801 and 6,880 gene families. Gene rarefaction analyses showed that the addition of future *E. anophelis* genomes to the analysis will still significantly increase the size of the pan-genome (12). The spectrum of gene frequencies for the *E. anophelis* pan-genome (Fig. 2b) showed that the vast majority of gene families were either encoded in a few genomes (39% in three or less) or in most of them (42% in more than 14 genomes), confirming that the genome sequencing of additional *Elizabethkingia* strains will uncover multiple novel gene families.

**FIG 2.**
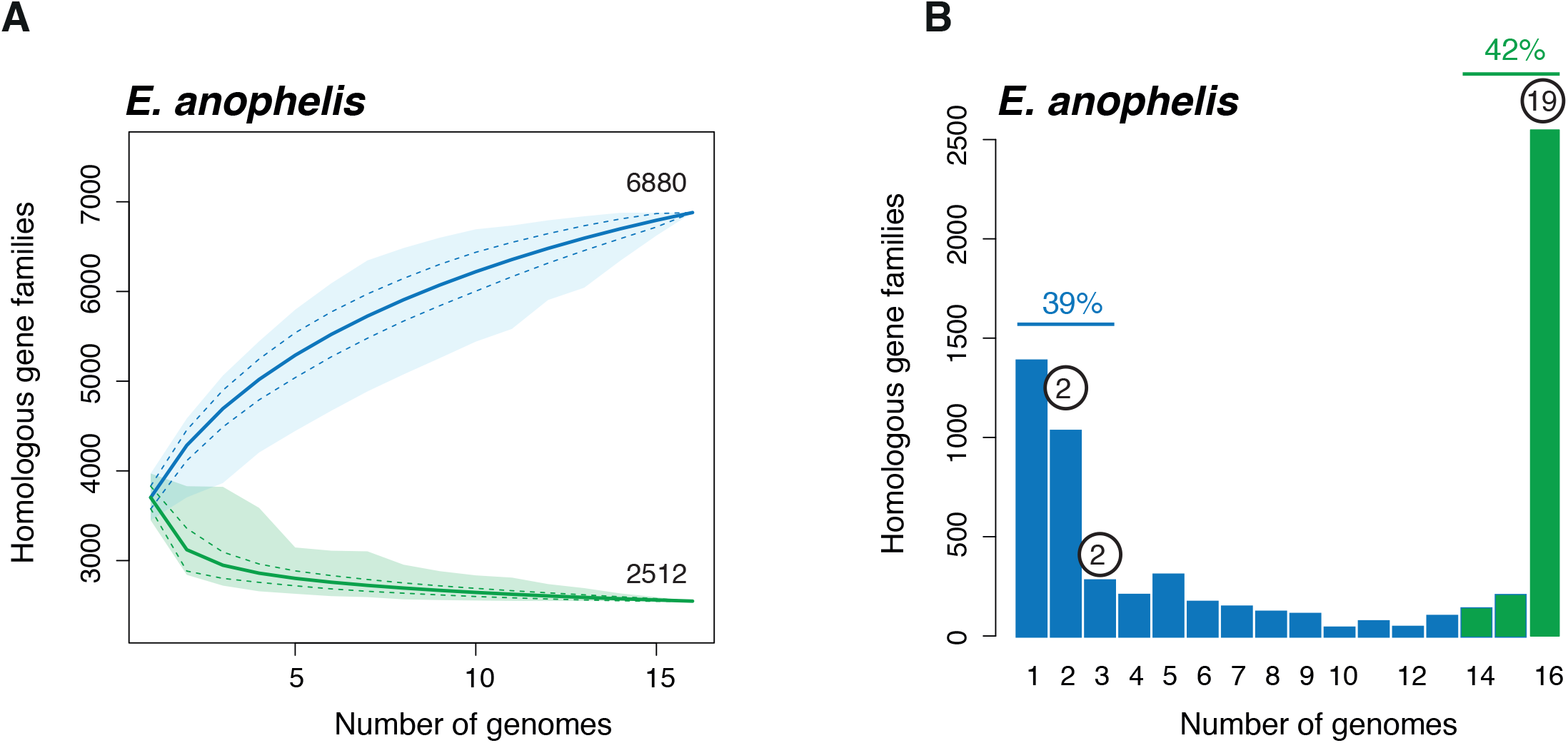
Core- and pan-genome sizes of *Elizabethkingia anophelis* (a) and spectrum of frequencies for *E. anophelis* gene repertoires (b). The pan- and core-genomes were used to perform gene accumulation curves. These curves describe the number of new genes (pan-genome) and genes in common (core-genome) obtained by adding a new genome to a previous set. The procedure was repeated 1,000 times by randomly modifying the order of integration of genomes in the analysis. The spectrum of frequencies represents the number of genomes where the families of the pan-genome can be found, from 1 for strain-specific genes to 16 for core genes. Blue indicates accessory genes and green the genes that are highly persistent in *E. anophelis*. Numbers in circles above the bars indicate the number of antimicrobial resistance (AR) gene families.

### Core genome-based phylogenetic structure of *E. anophelis*

To determine the phylogenetic origin of the African strains with respect to strains from other world regions, we selected a subset of 1,546 genes families with very reliable alignments (few indels) to infer robust phylogenies. Phylogenetic analysis based on concatenated alignments of the 1,546 gene sequences showed that all *E. anophelis* isolates were clearly separated from closely related species *E. miricola* (its sister group) and *E. meningoseptica* (Fig. 3a). These results fully corroborated the classification into three genomic groups based on ANI and 16S rRNA analyses.

**FIG 3.**
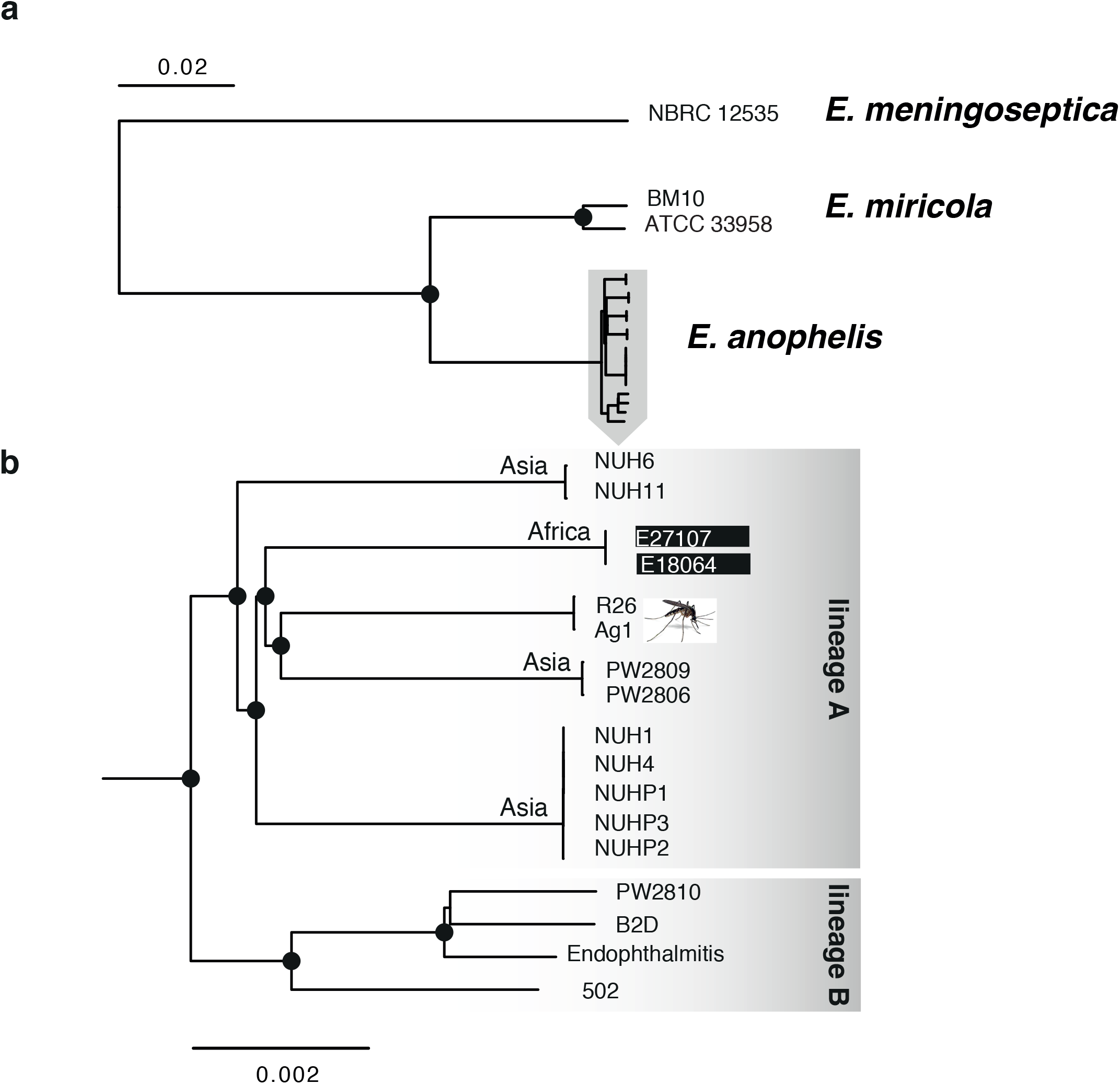
Phylogenetic tree of the *Elizabethkingia* genus based on the multiple sequence alignments of the 1,542 cgMLST loci. Black dots correspond to bootstrap supports higher than 90%.

The phylogenetic structure of *E. anophelis* revealed two main lineages, each containing distantly related sublineages (Fig. 3b). Whereas clinical isolates PW2810, 502, B2D and Endophthalmitis formed lineage B, all other human infection isolates were clustered into lineage A together with the mosquito strains Ag1 and R26. Therefore, *E. anophelis* mosquito strains share a common evolutionary history with a subset of *Elizabethkingia* isolates from humans. Nevertheless, the two mosquito strains were closely related and represented a distinct sublineage (named sublineage 1) that was separated by a large evolutionary distance from other isolates. Four additional sub-lineages were discerned within major lineage A, comprising respectively: i) Three patient isolates and two hospital environment isolates from a Singapore outbreak (7); ii) Two unrelated hospital environment isolates from the above outbreak investigation; iii) Strains PW2806 and PW2809 (HKU37 and HKU38, respectively) from Hong Kong (8); and iv) The two African strains from Bangui (6). Within each sublineage, the strains were highly related. These results show that the African strains formed a distinct sublineage, therefore having a clearly distinct evolutionary origin from Asian nosocomial strains and mosquito strains. The nearly simultaneous recognition of the emergence of *E. anophelis* infections in Africa and Asia thus reflects independent epidemiological events.

### Antimicrobial resistance-associated genomic features of African *E. anophelis* isolates

All *Elizabethkingia* genomes contained at least 17 antimicrobial resistance genes (Table S3). Most prominent among these were genes coding for beta-lactamases, including the metallo-beta-lactamase genes *bla_blaB_* and *bla_GOB_* (10, 13), and for efflux systems, which may contribute largely to the phenotypic resistance of *Elizabethkingia* isolates to most antibiotics. These findings are consistent with previous descriptions of multiple resistance genes in *Elizabethkingia* genomes (8, 11, 12, 14). Most resistance genes belonged to persistent families, i.e., they were part of the core genome of *Elizabethkingia*. This was the case for genes *bla_blaB_* and *bla_GOB_*, which were previously described as intrinsic and chromosome-borne in *E. meningoseptica* (10). We observed that both *bla_blaB_* and *bla_GOB_* were located in highly syntenic regions in all members of the *Elizabethkingia* genus, but were absent from the closely related genera *Flavobacterium, Riemerella, Chryseobacterium* (not shown). These results show that multiple potential resistance genes are ancestral in the genus *Elizabethkingia* and suggest acquisition by horizontal gene transfer into this genus before the evolutionary separation of the three *Elizabethkingia* species.

Interestingly, several additional resistance genes were detected in the African isolates (24 genes in total) as compared to other *Elizabethkingia* genomes (range: 17 – 20). Whereas all *Elizabethkingia* genomes contained one conserved chloramphenicol acetyltransferase (CAT) gene, the two African isolates contained an additional CAT gene that had 80.7% amino-acid identity with the conserved copy. The phylogenetic relationships of the conserved copy (Fig. S2) was concordant with the classification of the strains into *E. meningoseptica, E. miricola and E. anophelis*, indicating vertical evolution of the conserved CAT copy from an ancestral *Elizabethkingia* gene. In contrast, the additional CAT copy from African isolates was closely related to *Riemerella* sequences (Fig. S2), suggesting introduction by horizontal gene transfer from this genus.

Each of the African isolates also carried two additional metallo-beta-lactamases of the B1 (NDM-CcrA) subclass. In addition, an aminoglycoside acetyltransferase (AAC3-I) was found only in the two African isolates and in one isolate (NUH6) from Singapore. Finally, a *tet*(X) gene coding for a tetracycline inactivating enzyme 99.7% identical to that of *Bacteroides fragilis* (15) was found in the African isolates and in the Singapore isolate NUH4. Interestingly, the additional CAT gene, *tet*(X) and one subclass B1 beta-lactamase gene were clustered in a genomic region that also contained a putative class D beta-lactamase gene, a mercuric reductase gene and a lincosamide nucleotidyltransferase (LinF) gene associated with resistance to lincosamides (16). This region had high similarity to plasmid pRA0511 of *Riemella anatipestifer* (17). Furthermore, the other subclass B1 beta-lactamase gene and the AAC3-I gene were adjacent to each other and to a transposase gene. These observations indicate that the additional resistance genes of the African *E. anophelis* isolates were acquired through the transfer of mobile genetic elements.

### Identification and diversity of capsular polysaccharide synthesis clusters

Currently, the pathophysiology of *Elizabethkingia* infections is poorly understood (18). We searched for virulence-associated features in the African *E. anophelis* and other *Elizabethkingia* genomes by using the VFDB database (Table S4). A number of coding sequences showed homologs to iron and heme acquisition, to hemolysins and to oxidative stress resistance proteins, consistent with previous reports (8, 11, 12, 14). Among the hits found in the VFDB database, several corresponded to proteins putatively involved in capsule synthesis.

To our knowledge, *Elizabethkingia* strains were not previously reported as being capsulated or as harboring a *cps* cluster. Given that bacterial capsules represent one of the main bacterial virulence factors, particularly in pathogens that may be responsible for meningitis (19), we investigated whether the African isolates are capsulated. Using the India ink method, we observed a clearly distinguishable capsule, produced by the two strains during stationary phase when cultured either in BHI or in LB medium.

Further inspection of genome sequences revealed a Wzy-dependent *cps* gene cluster (20) comprising 27 co-oriented coding sequences in the two African isolates and in most other *Elizabethkingia* strains (Fig. 4). The upstream (3’) part of the gene cluster was highly conserved (Fig. 4) and included the two well-characterized genes *wzc and wza*. The former codes for an inner membrane protein necessary for capsular polymerization and translocation across the inner membrane. The latter encodes a channel that allows translocation of the polysaccharide across the outer membrane. Interestingly, these two genes were found in two copies in most strains separated by three other conserved genes (a RecX family transcription factor, and two other genes of unknown function).

**FIG 4.**
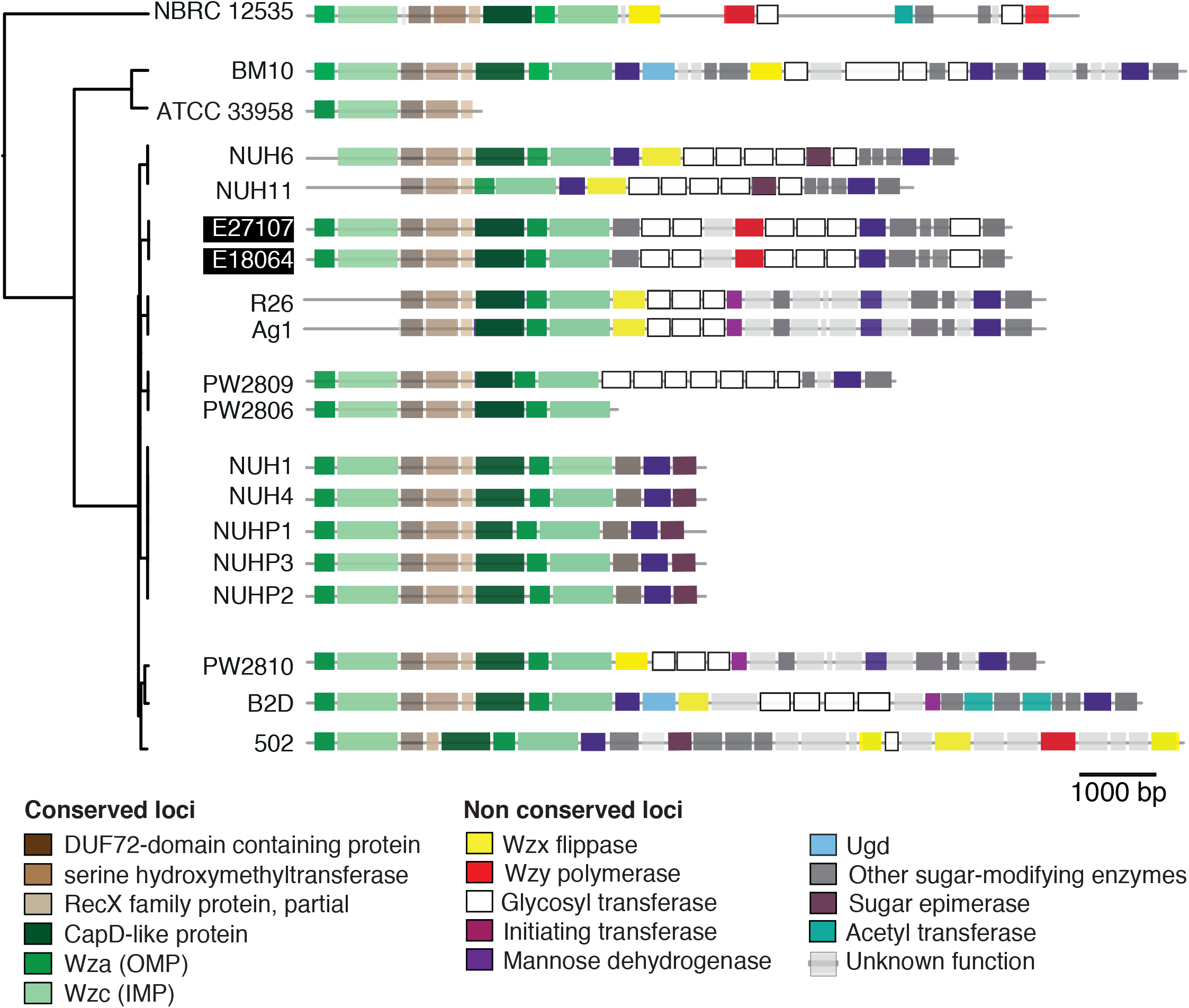
Genetic organization of the putative capsular polysaccharide synthesis (*cps*) gene clusters in *Elizabethkingia* genomes. Open reading frames (ORFs) are shown as rectangles and are drawn to scale. Intergenic spaces are not to scale. Three highly conserved genes, including *recX*, are depicted in brown. Green indicates highly conserved genes involved in capsule translocation (*capD, wza, wzc*). Yellow and red indicate genes *wzx* and *wzy*, respectively.

The downstream region was highly variable and may be responsible for the Wzy-dependent synthesis of polysaccharides with distinctive chemical compositions and which would possibly correspond to distinct capsular serotypes as observed in other capsulated species (21, 22). This region comprised the flippase (*wzx*) and the polymerase (*wzy*) and other genes coding for glycosyl transferases, acetyl transferases and other sugar-modifying enzymes. In some strains, the essential genes involved in polymerization - *wzx* and *wzy* - could not be confidently identified. These genes are known to be poorly conserved even among phylogenetically closely related isolates, rendering their detection difficult (23).

Interestingly, the genetic organization was strongly conserved among strains belonging to the same phylogenetic sublineage (as defined above; Fig. 4). These results suggest that capsular types are conserved within *E. anophelis* sublineages, but not between them. For example, the groups of African isolates, and mosquito isolates (R26 and Ag1) showed few intra-group and many between-group differences in the downstream regions of the capsular cluster locus. More specifically, the two African genomes seem to code for a unique and distinctive capsule, relative to other *Elizabethkingia* strains. Capsular type variation in *Elizabethkingia* may restrict immune cross-reactions and could be associated with distinct pathogenic properties, as observed in other capsulated bacterial groups (24-26). Overall, these results warrant future studies into the pathophysiological and epidemiological implications of capsular variation in *Elizabethkingia*.

### Secretion systems

Bacterial secretion systems are associated with important functions such as detoxification, antibiotic resistance and scavenging. We used MacSyFinder (27), together with TXSScan profiles (28), to identify the secretion systems encoded in the *Elizabethkingia* genomes (see Methods and **Table S5)**. While T9SS (PorSS) is present in many members of the phylum *Bacteroidetes* (29), we found none in the *Elizabethkingia* genomes. We also found no evidence for the presence of Type 2, 3, 5 protein secretion systems, which fits previous suggestions that these systems are absent from *Bacteriodetes* (28). In contrast, we found two families of T1SS in the genomes of *Elizabethkingia*, one of which was present in both African strains.

A third variant of the type VI secretion system (T6SSiii) was recently uncovered in Bacteroidetes (30). T6SS can be involved in bacterial competition (30) or pathogenicity (31). Interestingly, one locus encoding a T6SS^iii^ was found in every strain of our dataset. This system was highly conserved, except for the gene family tssI that showed extensive copy number variability. This gene encodes the cell-puncturing device and it can be fused with a variety of toxic domains, thus encoding a toxin (32). These systems might provide *Elizabethkingia* strains a way to antagonize competing bacteria in the complex environments such as the mosquito gut.

Type 4 secretion systems (T4SS) implicated in conjugative DNA transfer were previously described in *E. anophelis* (11, 14). We identified such systems in nearly all the genomes. Some strains encoded up to six conjugative systems (e.g., strain NUHP3), which is among the highest values observed among bacteria (33). Expectedly, all these systems were of the MPFB type, the one specific to Bacteroidetes. All the genomes encoding a T4SS also encoded one or several relaxases, suggesting that these systems are involved in conjugation of mobile genetic elements and not in protein secretion. The abundance of conjugative mobile genetic elements suggests that they might drive much of the genetic diversification observed in *Elizabethkingia*.

### Standardized strain typing method based on genome sequencing for *Elizabethkingia* epidemiology

In order to explore the epidemiological links between the African and other *E. anophelis* isolates, we translated whole genome sequences into high-resolution genotyping data. For this purpose, we followed the cgMLST strategy, which allows creating universal genotype nomenclatures and comparing isolates globally (34-36). A cgMLST scheme was developed based on 1,542 genes that could be aligned with high confidence among *Elizabethkingia* genomes. We then tested this scheme by scanning the 20 isolates for allelic variation at these loci. The identity of the 1,542 alleles was recorded for each isolate. The resulting allelic profiles were then clustered to obtain groups of isolates. These groups coincided with the sublineages from the phylogenetic analysis of the core gene sequences (Fig. S4).

To validate our cgMLST approach, we took advantage of the availability within the dataset of public genomes, of groups of isolates that were epidemiologically related, i.e., which were isolated during single outbreaks or were associated with direct mother-to-child transmission (7, 8). First, the five *E. anophelis* strains NUHP1, NUHP2, NUHP3, NUH1 and NUH4 from a single outbreak from Singapore were compared (7). We found that four of these strains were almost identical (2 to 6 cgMLST allelic mismatches), whereas NUH4 was slightly more distant (32 or more allelic mismatches with the others). This is highly consistent with the whole genome single nucleotide polymorphism (SNP) analysis by Teo and colleagues (12), who found 24 to 38 SNPs among the four closely related isolates, whereas at least 176 SNPs separated these from isolate NUH4. In a second study (8), two isolates (HKU37 and HKU38) from a mother and her neonate were considered to be directly related based on their genomic sequences, which showed no difference out of 2,000 genes. Based on our cgMLST approach, these two isolates were also identical, as none of the 1,542 loci was variable between them. In contrast, 1,433 cgMLST loci were distinct between this pair of isolates and a third independent isolate (HKU36) (8). These results demonstrate the extremely high efficiency of the cgMLST approach to distinguish epidemiologically related isolates from unrelated ones.

To our surprise, the two African *E. anophelis* isolates, which were isolated 5 years apart in the same hospital in Bangui (6), differed at only 4 alleles. This level of variation is similar to the variation observed among related isolates of the Singapore outbreak and thus strongly suggests to us that the two African hospital isolates may also be epidemiologically related. We therefore hypothesize that a single *E. anophelis* strain caused the two infections five years apart, due to its persistence in a hospital environmental source.

The core-genome MLST approach and attached genomic sequences were made publicly available through the Institut Pasteur server (http://bigsdb.web.pasteur.fr/elizabethkingia/).

### Clustered regularly interspaced short palindromic repeats (CRISPR)

CRISPR loci exhibit spacer diversity that reflects a history of prior invasions by different phages and plasmids (37). In addition, the high variability of CRISPR loci in some bacterial species implies that they represent useful markers for epidemiological investigations (38, 39). To extend our understanding of genomic variation between the African isolates and among the other *Elizabethkingia* isolates, we investigated CRISPR-Cas systems and analyzed spacer content in the 20 strains. All three species contained subtype II-C CRISPR-Cas systems comprising of Cas1, Cas2, and Cas9 proteins, while no homolog of Cas4 and Csn2 were identified (40). The loci were characterized by unusually long repeats (that is, 47 bp in size, Fig. 5a), as previously observed in bacteria belonging to the *Bacteroidetes* phylum (40). However, the CRISPR locus was only present in a few *Elizabethkingia* strains (Fig. S3). These results demonstrate the occurrence of CRISPR systems in *Elizabethkingia* and the variable presence of these systems among phylogenetic sublineages of *E. anophelis*.

**FIG 5.**
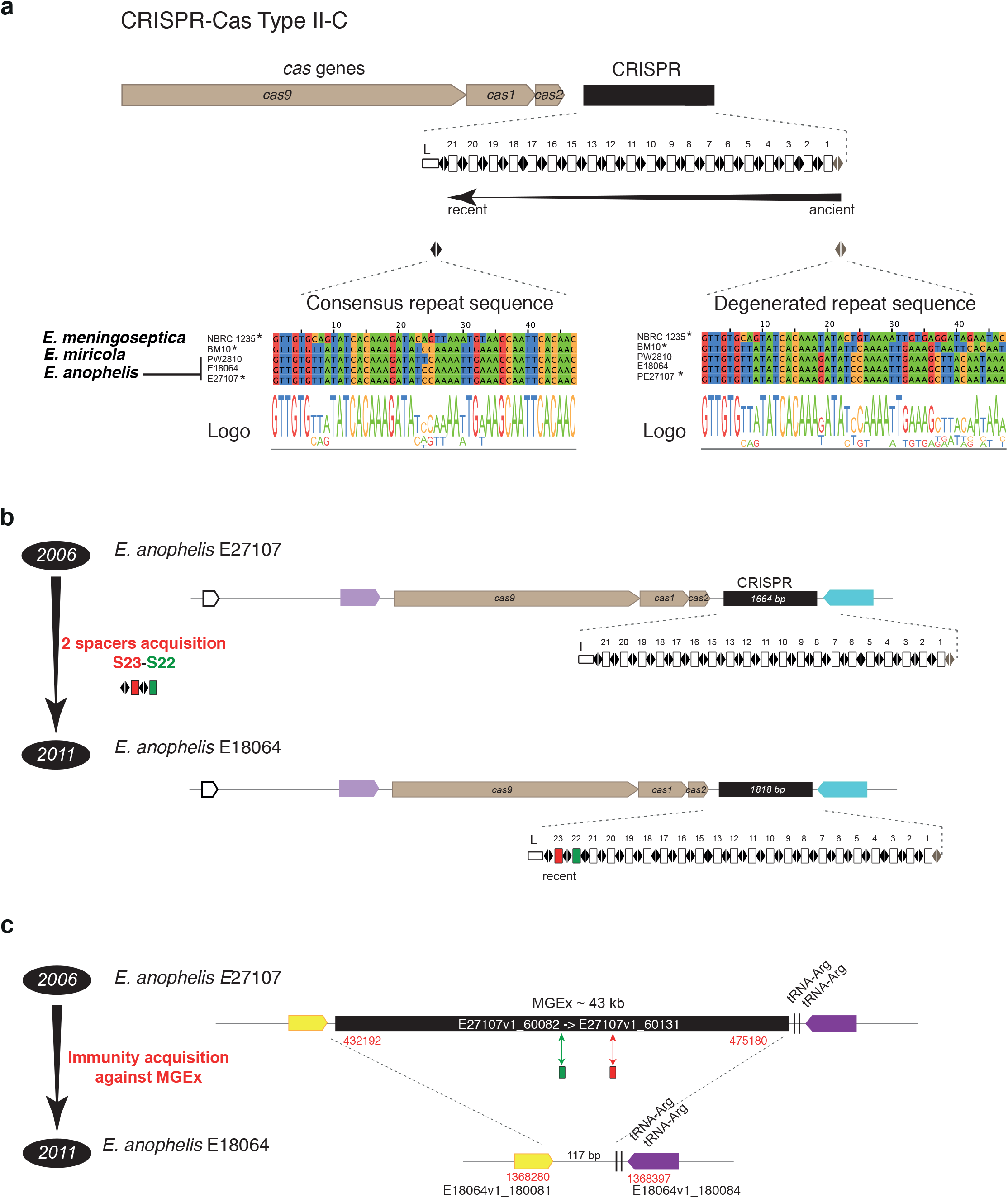
CRISPR-Cas type II-C organization and diversity. (a) System organization is depicted on the top, with *cas* genes in brown, and the repeat-spacer array in black. Below the gene scheme, the repeat and spacer (captured MGEs) content is detailed as black diamonds and white rectangles, respectively. L corresponds to the leader sequence. Bottom line, consensus repeat sequence and degenerated repeat sequence in the 5 genomes containing such system. (b) Diversity of spacer content between the two Central African Republic hospital isolates. (c) Presence of a large MGE containing perfect matches with the two additional spacers S22 and S23.

Analysis of the spacer diversity among *Elizabethkingia* CRISPR arrays revealed no shared spacer between or within species, with the exception of the two hospital acquired African isolates, which showed a strong CRISPR conservation (Fig. 5b). This finding suggests a high rate of gain and loss of complete CRISPR-Cas system during *E. anophelis* evolution. CRISPR arrays of the two African isolates were almost identical, with the exception that E18064, which was isolated 5 years later (in 2011), contained two additional spacers at the 5’ end of the array (called S22 and S23, Fig. 5b). Importantly, according to the current spacer acquisition model, these two spacers are the most recent acquired spacers (see Methods). This clear genetic difference between CRISPR arrays of the African isolates was confirmed by PCR. CRISPR-based typing may thus offer the high level of discrimination needed for strain subtyping in a local epidemiological context.

Surprisingly, while the S22 and S23 spacers were clearly absent in the CRISPR array of the earlier African isolate E27107 (isolated in 2006), we found perfect matches (*i.e*. protospacers) in one E27107 specific genomic region of 43 kb (not including the CRISPR array) (Fig. 5c). This particular genomic island was totally absent in other strains and was characterized by the presence of a putative integrase gene located near a tRNA-Arg gene, three genes encoding putative phage tail and lysis proteins, with the remaining genes having unknown functions. Further, this region contained only co-oriented genes. Thus, we hypothesize that this putative mobile genetic element (MGEx) encodes a prophage. In sum, whereas isolate E27107 contained a shorter CRISPR array devoid of S22 and S23 and a putative prophage harboring S22 and S23 protospacers, isolate E18064 harbored the two spacers in its CRISPR array and lacked the corresponding prophage. Based on these observations, we speculate that the presence of these two spacers provides resistance against this particular MGE. The CRISPR and phage dynamics uncovered here emphasize the rapid diversification of MGE-related genomic features in *Elizabethkingia* genomes, even at the short evolutionary timescale of 5 years that separates the two African hospital infections.

## CONCLUSIONS

*Elizabethkingia anophelis* was recently recognized as a cause of nosocomial infections, and more recently, of community-acquired infections. In this work, the genome sequences of two hospital meningitis African isolates were established and compared with available genomes from other regions and sources. Our comparative phylogenetic analyses led us to refine the identification of *Elizabethkingia* strains and to demonstrate firmly the phylogenetic distinctness of the three described species of the genus. Comparative genomics analysis revealed specific features of the African isolates including additional resistance genes, a unique CRISPR locus and a specific putative capsular synthesis (*cps*) cluster. Demonstration of the capsulated nature of the African *E. anophelis* isolates, and the presence of variable *cps* clusters suggests an important determinant of the pathogenicity mechanisms and virulence heterogeneity among *Elizabethkingia* strains. Analysis of the phylogenetic structure of *E. anophelis* revealed several well-demarcated sublineages and demonstrated a distinct evolutionary origin of African clinical isolates, excluding an epidemiological link between recent African and Asian hospital infections. The core-genome MLST approach defined here and made publicly available through the Institut Pasteur bacterial genome database and analysis platform (http://bigsdb.web.pasteur.fr/elizabethkingia/) represents a standardized high-resolution genotyping tool. It will enable to share a common language on *E. anophelis* isolates sublineages, which will facilitate collaborative work on the population biology, epidemiology and pathophysiology of this newly recognized bacterial species. Here, the analysis of nosocomial infection isolates using this genotyping approach suggested that the hospital environment, rather than mosquitoes, acts as a reservoir for *E. anophelis*, and that the high persistence of the pathogen in this environment, if uncontrolled, allows for long-term transmissions.

## MATERIALS AND METHODS

### Isolates for genome sequencing

The two clinical strains E18064 (alias: V0378064) and E27107 (alias: Po0527107) isolated in Central African Republic were sequenced with a 100 base pair (bp) paired-end protocol in Illumina HiSeq-2000. Libraries were constructed with the Nextera DNA Sample Prep Kit (Illumina). Genome assembly was performed with the CLC Assembly Cell analysis package version 3.2.2. The average number of contigs and the N50 statistic values (i.e., the length for which half of the bases of a draft genome are situated in contigs of that length or longer) observed from the two assembled genomes were similar to those of the publicly available draft genomes (Table S1). The draft genome sequences of strains E18064 and E27107 were annotated using the MicroScope/MaGe platform (41) and were deposited in the European Nucleotide Archive under accession numbers CCAB000000000 and CCAC000000000, respectively.

### Genomes from sequence database

The eighteen *Elizabethkingia* genomic sequences available on April 1^st^, 2015 were downloaded from the NCBI genome sequence repository (www.ncbi.nlm.nih.gov/genome). The sequences comprised one complete genome and 17 whole-genome shotgun sequences available as scaffolds or contigs. The strains were initially identified as *E. anophelis* (12 isolates)*, E. meningoseptica* (n=4), *E. miricola* (n=1), and *Elizabethkingia* sp. (n=1) (Table S1).

### Average nucleotide identity and 16S sequence analysis

Average nucleotide identity values were estimated from the genomic assemblies using JSpecies v1.2.1 with the BLASTN option (42). 16S rRNA sequences were gathered from genomic sequences by BLASTN similarity search using the 16S sequence of the taxonomic type strain R26^T^ (GenBank accession number EF426425) as a query.

### Core genome and pan-genome

Core genomes (the set of genes present in all isolates) were built either for the genus or for *E. anophelis* species. Orthologs were identified as bidirectional best BLASTP hits (43) between the proteome of *E. anophelis* NUHP1 as a pivot and each of the other proteomes (18 for the genus and 15 for the species). Strain Endophthalmitis contained a large fraction of pseudogenes (likely due to low quality of the sequence) and was therefore excluded from this analysis. Hits with less than 40% (genus) or 80% (species) amino acid sequence similarity or more than 20% difference in sequence length were discarded. Genomes from the same species typically show low levels of genome rearrangements and this information can be used to identify orthologs more accurately (44, 45). Therefore, the core-genome of the species was defined as the intersection of pairwise lists of strict positional orthologs (as in (46)). The core-genomes consist in the genes present in all genomes of the two sets. They were defined as the intersection of the lists of orthologs between pairs of genomes.

Pan-genomes were built by clustering homologous protein-coding sequences (CDS) into families. We determined the lists of putative homologs between pairs of genomes with BLASTP used the E-values (<10^−4^) to perform single-linkage clustering with SiLiX v1.2 (47). A CDS is thus included in a family if it shares a relation of homology to at least one CDS already belonging to the family. SiLiX parameters were set to consider two CDS as homologs if the aligned part showed at least 60% (Elizabethkingia genus) or 80% (E. anophelis) identity and represented more than 80% of the smallest CDS length. The pan-genomes thus represent the full complements of genes in the genus and in the species, respectively. The pan-genomes of Elizabethkingia and of E. anophelis were determined independently.

### Core genome multilocus sequence typing (MLST)

To obtain a subset of CDSs that are highly reliable as genotypic markers, a subset of the core genome was selected based on length conservation criterion leading to the selection of 1,546 CDSs. These loci together constitute a cgMLST scheme useful for genotyping of *E. meningoseptica, E. anophelis*, and *E. miricola* isolates. The scheme comprises fewer loci than the core genome as expected from our more stringent selection criteria. The genome sequences were then scanned for allelic variation using the BIGSdb tool (35) using the allele sequences from reference strain R26 as the initial query sequences.

### Characterization of gene associated with drug resistance and horizontal gene transfer mechanisms

Acquired antimicrobial resistance (AR) genes were detected using HMMER3 (48) against the ResFams, a curated database of AR protein families and associated profile hidden Markov models, with the tblout and cut_ga options (49). Virulence-associated genes were searched on the VFDB (50) using BLASTP (minimum 40% identity with E-value <10-5), as in (14).

#### Comparison of genomes

Local genomic alignments visualization was performed using software BioNumerics v.7.5 (Applied-Maths, Sint-Martens Latem, Belgium) with default seed matching and stretch extension parameters.

### Phylogenetic analyses

For each of the 1,546 cgMLST loci, a multiple amino acid sequence alignment was performed with MAFFT v.7.205 (default options) (51) that was back-translated in order to obtain a codon-level alignment. The concatenation of these 1,546 multiple sequence alignments was used to estimate the p-distance between each pair of isolates. These estimated evolutionary distances were used to infer a genome-level phylogenetic tree with FastME v.2.07 (Balanced Minimum Evolution criterion, SPR-based BME tree search) (52).

Phylogenetic tree analyses from single markers (i.e. 16S rRNA and CAT) were performed from well-suited characters selected by BMGE v.1.12 (up to 50% allowed gap proportion with models PAM1 and BLOSUM62, respectively) (53) from the multiple sequence alignments generated by MAFFT. Tree inferences from 16S rRNA and CAT sequences were performed by PhyML v.20131016 (54) (SPR-based ML tree search with evolutionary models GTR+I and LG+Γ4+I, respectively).

### Detection of CRISPR-Cas systems

Clusters of *cas* genes were identified and classified using MacSyFinder (27). To identify *cas* pseudogenes, all Cas protein sequences previously detected were searched in all the *Elizabethkingia* draft genomes using TBLASTN (E-value <10-3). CRISPR arrays were identified following a previously published methodology (55). In short, they were identified using CRT (CRISPR Recognition Tool) with the option -maxRL=60 (56), in all the Elizabethkingia draft genomes. For each array, the repeats were extracted and were aligned using MUSCLE (57). Then, we used cons (www.bioinformatics.nl/cgi-bin/emboss/help/cons) to obtain consensus sequences from these multiple sequence alignments of each array. In all cases, the consensus sequence corresponds to the most frequent sequence within a particular array. We used the consensus sequence of the repeats as patterns to identify additional, smaller and/or degenerate repeat clusters in all draft genomes with fuzznuc (www.bioinformatics.nl/cgi-bin/emboss/help/fuzznuc). This step has also confirmed the lack of any repeat in some draft genomes devoid of detectable CRISPR array and cas genes.

### Spacer diversity

To identify additional spacers (i.e located at contig extremities) or to verify the lack of a particular spacer in a given strain, all spacer sequences previously detected (i.e., flanked by repeats at both sides) were searched in all the *Elizabethkingia* draft genomes using BLASTN. Only hits showing at least 90% identity with the query, less than 20% difference in sequence length and flanked by sequences similar to repeats were considered as additional spacers. We then compared the spacer content within and between strains and species. Therefore, two spacers were considered as similar (within strain) or common (between strains) if they had less than 20% difference in sequence length and at least 90% identity.

### CRISPR locus orientation for polarized spacer acquisition

During the adaptation phase, spacer acquisition occurs in a polarized fashion: new spacers are typically integrated at the 5’ (leader) end of the CRISPR array, which involves the duplication of the first repeat of the array, and older spacers and repeats decay at the 3′ end. Consistent with this spacer acquisition model, we found that the most degenerated repeat was located at one end of each CRISPR arrays. Hence, such particular repeat was considered as the signature of the 3’ end of the array, which corresponds to the oldest part of the locus. Such positional information represents a timeline of spacer acquisition events.

#### CRISPR locus PCR

To confirm the genomic differences observed between CRISPR arrays of the two African isolates based on Illumina sequence data, three primer pairs were designed for PCR amplification and sequencing of the CRISPR locus in three overlapping fragments: (*i*) L1F- CGT AAA CGT CTG TTA GAT GAT GG and sp15R-AAA CCA TTC TAC GGA GAA C; (*ii*) sp17F-GAT GTA ATA AGA GTT GTT GCG and sp5R-TCG GAT TTA TGA GGT GAT CCC; and (*iii*) sp7F-CAT AGA TCA CAC ATA CAG GGC and L1R-TGA GCG CCC ATG TTG TCT CCG. PCR conditions for all amplification reactions were as follows: initial denaturation at 94°C for 5 min; 30 cycles at 94°C for 30 s, 50°C for 30 s, and 72°C for 30s; and final extension at 72°C for 5 min. PCR products were purified by ultrafiltration (Millipore), and nucleotide sequences were obtained using the PCR primers and BigDye Terminator v1.1 chemistry (Applied Biosystems, Foster City, CA) on an ABI 3730XL apparatus (Applied Biosystems, Foster City, CA). Sequence traces were edited and assembled using BioNumerics.

### Detection of capsular gene clusters

To identify capsular gene clusters, we performed a keyword search of the Pfam database (pfam.xfam.org) for protein profiles involved in capsular polysaccharide production such as glycosyl transferases, ABC transporters, *wzx* flippase and *wzy* polymerase (Table S6). We then performed a search of these profiles in *Elizabethkingia* genomes using HMMER3 (48), with the tbl_out option. This allowed us to clearly identify two capsule clusters in each of the strains 502 and NBRC 12535. These clusters were both located in the same genomic region, which included a highly conserved RecX family transcriptional regulator. We then searched for *recX* in the other genomes and reconstructed the putative capsular polysaccharide synthesis (*cps*) cluster in the *recX* neighborhood using the aforementioned protein profiles.

## ACKNOWLEDGMENTS

We would like to express our most sincere thanks to the director of Institut Pasteur in Bangui, Dr Mirdad Kazanji, who made this work possible.

## Supplementary tables

**TABLE S1**. General characteristics of the 20 *Elizabethkingia* genomes analyzed in this study.

**TABLE S2**. Average nucleotide identity (58) values computed from 20 *Elizabethkingia* genome sequences.

**TABLE S3**. Antimicrobial resistance-associated features identified in *Elizabethkingia* genomes.

**TABLE S4**. Predicted virulence factors identified in *Elizabethkingia* genomes.

**TABLE S5**. Protein secretion systems identified in *Elizabethkingia* genomes.

**TABLE S6**. Proteins families searched in *Elizabethkingia* genomes to characterize capsular polysaccharide synthesis clusters.

## Supplementary figures

**FIG S1**. Phylogenetic analysis of *Elizabethkingia* isolates based on 16S rRNA sequences. a. Phylogenetic tree with bootstrap-based support (500 replicates) >70% indicated at branches. b. Detail of the multiple 16S rRNA sequence alignment (aligned characters 963-1,033).

**FIG S2**. Gene tree of chloramphenicol acetyltransferase (CAT) genes. Blue: sequences from Schwarz et al. 2004, FEMS Microbiology Reviews 28:519–542; magenta: sequences from *Elizabethkingia* strains; black: closely related homologous sequences selected from RefSeq. NCBI accession ids are given on the right. Bootstrap-based supports (500 replicates) >70% are indicated at branches.

**FIG S3**. Distribution of CRISPR-Cas systems in *Elizabethkingia* genus. White indicates absence of the trait and black indicates its presence. The numbers reported on the right correspond to the number of spacers identified in each genome.

**FIG S4**. Gene-by-gene comparison of the *Elizabethkingia* genomes based on 1,542 loci included in the core genome multilocus sequence typing (cgMLST) scheme. a: UPGMA dendrogram computed from the matrix of allelic mismatches distances (i.e., the proportion of distinct alleles among each pair of profiles). b: Minimum spanning tree of the Singapore lineage. The number of allelic mismatches between each displayed link is given. c: Minimum spanning tree of the African lineage, which differ by 4 alleles out of 1,542.

